# Mechanism of nucleic-acid-driven LLPS of TDP-43 PLD

**DOI:** 10.1101/2022.11.01.514785

**Authors:** Dang Mei, Tongyang Li, Shibo Zhou, Jianxing Song

## Abstract

Most membrane-less organelles (MLOs) formed by LLPS contain both nucleic acids and IDR-rich proteins. Currently while IDRs are well-recognized to drive LLPS, nucleic acids are thought to exert non-specific electrostatic/salt effects. TDP-43 functions by binding RNA/ssDNA and its LLPS was characterized without nucleic acids to be driven mainly by PLD-oligomerization, which may further transit into aggregation characteristic of various neurodegenerative diseases. Here by NMR, we discovered unexpectedly for TDP-43 PLD: 1) ssDNAs drive and then dissolve LLPS by multivalently and specifically binding Arg/Lys. 2) LLPS is driven by nucleic-acid-binding coupled with PLD-oligomerization. 3) ATP and nucleic acids universally interplay in modulating LLPS by competing for binding Arg/Lys. However, the unique hydrophobic region within PLD renders LLPS to exaggerate into aggregation. The study not only unveils the first residue-resolution mechanism of the nucleic-acid-driven LLPS of TDP-43 PLD, but also decodes a general principle that not just TDP-43 PLD, all Arg/Lys-containing IDRs are cryptic nucleic-acid-binding domains that may phase separate upon binding nucleic acids. Strikingly, ATP shares a common mechanism with nucleic acids in binding IDRs, thus emerging as a universal mediator for interactions between IDRs and nucleic acids, which may underlie previously-unrecognized roles of ATP at mM in physiology and pathology.

## Introduction

Liquid-liquid phase separation (LLPS) has been recently established as the common principle for forming membrane-less organelles (MLOs), which include nucleoli, Cajal bodies, nuclear speckles, paraspeckles, histone–locus bodies, nuclear gems, and promyelocytic leukemia (PML) bodies in the nucleus as well as P-bodies, stress granules, and germ granules in the cytoplasm (1-5). Intriguingly, most, if not all, MLOs are composed of both nucleic acids and proteins rich in intrinsically-disordered regions (IDRs). Currently, while IDRs have been well-recognized to drive LLPS, the effect and mechanism of nucleic acids to affect LLPS remain largely elusive. In particular, one fundamental question remaining to be answered is whether nucleic acids affect LLPS just by non-specific electrostatic/salt effects or by specific binding. As a consequence, the high-resolution mechanism is completely absent for nucleic acids to interact with IDRs in LLPS.

TAR-DNA-binding protein-43 (TDP-43) was originally identified to be a human protein capable of binding the trans-active response (TAR) DNA region of HIV to repress its transcription (6). Now TDP-43 has been uncovered to be a nuclear protein which functions by binding a large array of RNA and single-stranded DNA (ssDNA) including more than 6000 RNA species of diverse sequences (7,8). Furthermore, in response to stresses, TDP-43 also becomes localized into the cytoplasm to participate in forming stress granules (SGs) with RNAs through phase separation. Most intriguingly, under pathological conditions, TDP-43 accumulates and forms inclusion bodies in the cytoplasm of motor neurons, which is a common pathological hallmark of most cases of amyotrophic lateral sclerosis (ALS), as well as associated with an increasing spectrum of other major neurodegenerative diseases, including Alzheimer’s (AD), Parkinson’s (PD), frontotemporal dementia (FTD), and Huntington’s (HD) diseases (7-14).

The 414-residue TDP-43 is composed of the folded N-terminal domain (NTD) (15,16) and two RNA recognition motif (RRM) domains for binding various RNA and DNA (17-20), as well as the C-terminal low-complexity (LC) domain over the residues 265-414 with the amino acid composition similar to those of yeast prion proteins, thus designated as the prion-like domain (PLD) (14,21,22). Puzzlingly, although TDP-43 PLD is intrinsically disordered, it hosts almost all ALS-causing mutations identified so far (7-9). Intriguingly, as compared with PLDs in other RRM-containing proteins, TDP-43 PLD is unique in owning a hydrophobic region over residues 311-343, which adopts the partially-folded helical conformation in solution (21,22). Recently, TDP-43 PLD has been characterized to critically drive LLPS of TDP-43, which is essential for forming SGs but might further exaggerate into pathological aggregates or amyloid fibrils (21-31). So far, LLPS of TDP-43 PLD has been extensively characterized without nucleic acids to be driven mainly by the formation of the dimeric/oligomeric helix over the unique hydrophobic region (22,25,26), which appears also to undergo conformational exchanges with the amyloid-like β-rich oligomers (30).

Nevertheless, so far it remains unknown how nucleic acids modulate LLPS of TDP-43 PLD, non-specifically or specifically? although cellular environments where TDP-43 plays its physiological and pathological roles are rich in various nucleic acids and LLPS of TDP-43 has been indeed found to be affected by both RNA (32) and ssDNA (33). Furthermore, ATP, the universal energy currency mysteriously with concentrations (>mM) much higher than required for its classic functions in all living cells (34), was shown to act as a biological hydrotrope to dissolve LLPS of RRM-containing proteins including TDP-43 at >5 mM (35,36). Very recently, we found that ATP is capable of biphasically modulating LLPS of TDP-43 PLD: namely induction at low but dissolution at high ATP concentrations (37). Most unexpectedly, despite containing only five Arg and one Lys residues within the 150-residue PLD: namely Arg268, Arg272, Arg275, Arg293 and Arg361, as well as Lys408, our residue-specific NMR results unambiguously indicate that ATP achieves the biphasic modulation of LLPS by acting as a bivalent binder to specifically binding to Arg/Lys residues through electrostatic interactions between ATP triphosphate group and Arg/Lys side chain cations as well as π-π/π-cation interactions between ATP purine ring and Arg/Lys side chains (37). Interestingly, the binding affinity of ATP to Arg was experimentally shown to be much higher than that to Lys (37-39). Here as assisted by turbidity measurement and DIC imaging, we utilized NMR spectroscopy to visualize the effects on LLPS and binding events of Tar32, a natural ssDNA ligand of TDP-43 together with two non-specific ssDNAs A6 and A32 with the wild-type (WT) and two mutated PLDs: namely Del-PLD with residues 311-343 deleted and AllK-PLD with all five Arg replaced by Lys residues (37). Although ssDNA and RNA only has two minor differences in chemical structures: the base thymine (T) in DNA is replaced by uracil (U) in RNA, while d-2-deoxyribose in DNA is replaced by d-ribose in RNA, unlike RNA which is vulnerable to degradation by RNAse extensively existing in environments, ssDNA has a very high chemical stability, which thus allows to acquire time-consuming NMR spectra.

The results decode for the first time: 1) all three ssDNAs could drive and then dissolve LLPS of TDP-43 WT-PLD with the capacity dependent on the length but not sequence. Most unexpectedly, ssDNAs modulate LLPS by specifically binding Arg/Lys with the affinity to Arg higher than that to Lys. 2) In the presence of nucleic acids, LLPS of TDP-43 PLD is driven by the nucleic-acid-binding coupled with PLD-oligomerization. Nevertheless, in the general context, the multivalent nucleic-acid-binding itself is sufficient to drive LLPS of Arg/Lys-containing IDRs without the need of any other driving force. 3) ATP and ssDNAs interplay in modulating LLPS by competing for binding Arg/Lys. However, for TDP-43 PLD, the presence of the unique hydrophobic region renders its LLPS to be prone to exaggerating into aggregation. The study unveils the first residue-resolution mechanism of the nucleic-acid-driven LLPS of TDP-43 PLD, which emphasizes the specific role of the multivalent nucleic-acid-binding in driving LLPS as well as the unique potential of the hydrophobic region to exaggerate functional LLPS into pathological aggregation. Remarkably, a general principle is emerging that not just TDP-43 PLD, all Arg/Lys-containing IDRs, which account for a large portion of eukaryotic proteomes, are cryptic nucleic-acid-binding domains and may undergo LLPS upon multivalently binding nucleic acids. Most strikingly, ATP shares a common mechanism with nucleic acids in binding Arg/Lys residues of IDRs, thus emerging as a general mediator for interactions between IDRs and nucleic acids, which may represent a key mechanism underlying previously-unrecognized roles of ATP at mM in physiology and pathology.

## Results

### ssDNAs biphasically modulate LLPS with the length-dependent capacity

Previously, by NMR we have characterized the solution conformation and amyloid fibrillation (21) as well as its interaction with ATP (37) for the 150-residue TDP-43 PLD over residue 265-414 with a pI of 10.78 (Fig. 1a). Here by measurement of turbidity (absorption at 600 nm) and DIC imaging at the same protein concentrations (15 μM) and in the same buffer which we previously used (37), we first assessed how LLPS of TDP-43 WT PLD is affected by a 32-mer ssDNA Tar32, a natural ligand of TDP-43, whose sequence is derived from the trans-active response (TAR) DNA of HIV (6,33), and contains all four nitrogenous bases (Supplementary Figure 1).

**Fig 1.**
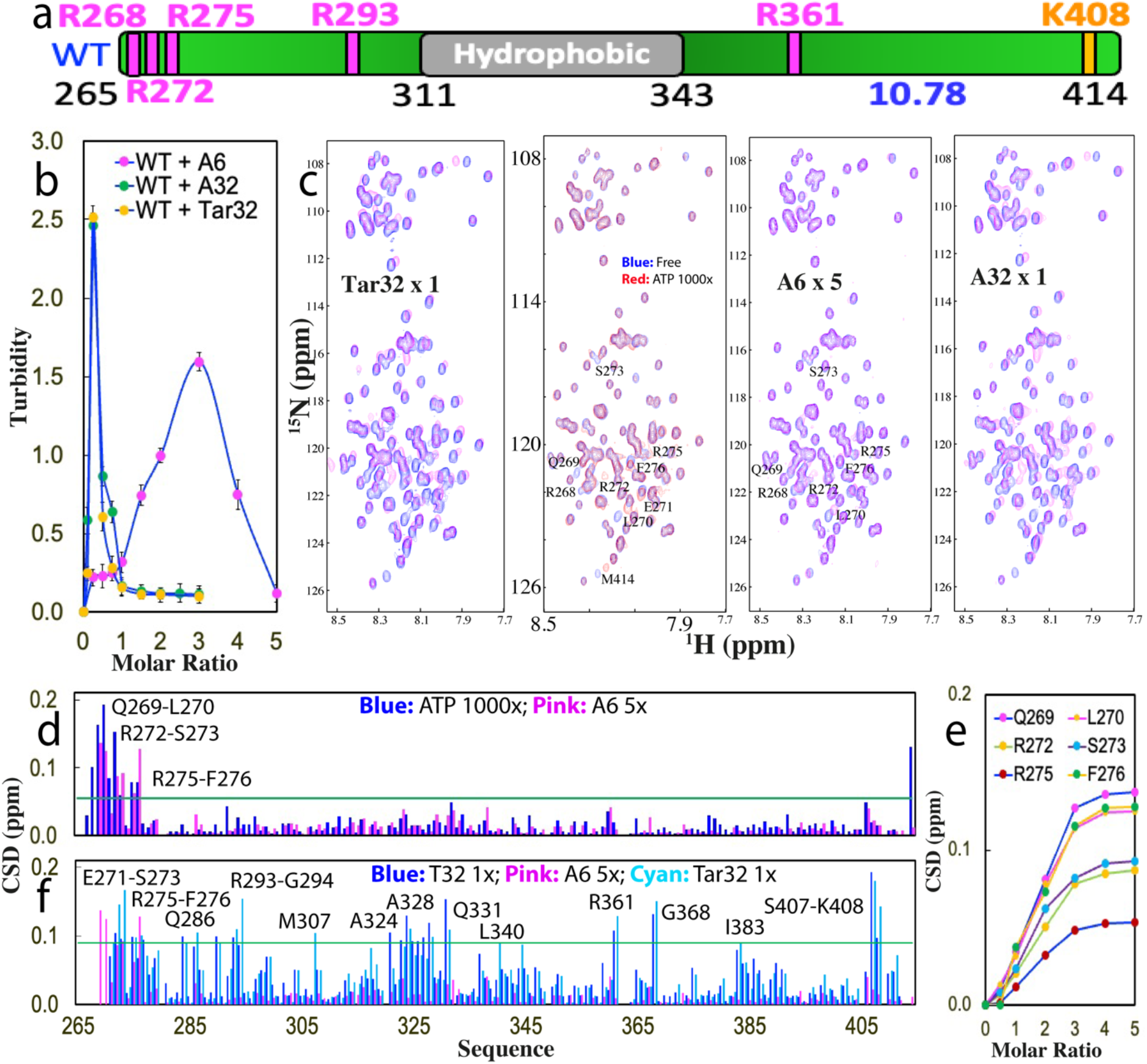
Three ssDNAs length-dependently modulate LLPS of TDP-43 WT-PLD. (a) Schematic representation of TDP-43 WT-PLD with five Arg and one Lys, as well as pI value indicated. (b) Turbidity curves (absorption at 600 nm) of WT-PLD in the presence of Tar32, A6 and A32 at different molar ratios. (c) ^1^H-^15^N NMR HSQC spectra of the ^15^N-labeled WT-PLD in the absence (blue) and in the presence of ATP at a molar ratio of 1:1000 (red), as well as A6 at 1:5, A32 and Tar32 respectively at 1:1 (purple). (d) Chemical shift difference (CSD) of WT-PLD between the free state and in the presence of ATP at 1:1000 (blue) or A6 at 1:5 (purple) respectively. The green line is used to indicate the value (0.05): Average + STD at the ratio of 1:5. The residues with CSD values > Average + SD are defined as significantly perturbed residues and labeled. (e) CSD tracings of six significantly perturbed residues in the presence of A6 at different ratios. (f) Chemical shift difference (CSD) of WT-PLD between the free state and in the presence A6 at 1:5 (purple), A32 at 1:1 (blue) and Tar32 at 1:1 (cyan) respectively. The green line is used to indicate the value (0.09): Average + STD in the presence of Tar32 at the ratio of 1:1 (PLD: Tar32). The residues with CSD values > Average + SD are defined as significantly perturbed residues and labeled.

Strikingly, upon stepwise addition of Tar32, the turbidity of TDP-43 PLD rapidly increased with a value of 0.25 at a molar ratio of 1:0.1 (PLD:Tar32), and then reached the highest value of 2.51 at 1:0.25 (Fig. 1b). DIC imaging showed that many liquid droplets were formed, some of which have diameters of ∼2 μm (Supplementary Figure 2). Intriguingly, further addition of Tar32 led to the reduction of turbidity and at 1:1, the turbidity value was only ∼0.11, where no droplet could be detected by DIC imaging. The results clearly indicate that Tar32 is able to biphasically modulate LLPS of TDP-43 WT PLD: namely to drive LLPS at low Tar32 concentrations but dissolve at high concentrations.

To assess whether the unique sequence of Tar32 is essential for modulating LLPS, we subsequently used a 32-mer ssDNA A32 containing only adenine (A) (Supplementary Figure 1) to titrate WT PLD under the same conditions. As shown in Fig 1b, the turbidity curve of A32 is very similar to that of Tar32 with the highest vale at 1:0.25. DIC visualization indicates that A32 also drove and then dissolved the formation of droplets with the overall pattern very similar to that of Tar32. Therefore, this set of results suggests that the specific sequence of Tar32 is not essential for biphasically modulating LLPS.

Next, we set to test whether the length of ssDNA is critical by titrating WT-PLD with a 6-mer ssDNA A6. As shown in Fig 1b, for A6, the turbidity reached the highest (1.59) at 1:3 and at 1:5 the turbidity reduced to 0.12 (Fig 1b). DIC imaging indicates that A6 did drive and then dissolve the formation of the droplets, but the number induced by A6 at 1:3 is less than that by Tar32 or A32 at 1:0.25, thus resulting in a smaller value. The results with Tar32, A32 and A6 together suggest that it is the length of ssDNA but not the sequence which is the key determinant for the capacity of ssDNA in biphasically modulating LLPS of TDP-43 PLD.

### NMR visualization of the biphasic modulation of LLPS by ssDNAs

To insight into the binding events for ssDNAs to drive and subsequently dissolve LLPS of TDP-43 WT-PLD, under exactly the same conditions as above for turbidity measurement and DIC imaging, we monitored the titrations of Tar32, A6 and A32 into WT-PLD by NMR HSQC spectroscopy, which is particularly powerful in pinpointing the binding events with a wide range of affinity at residue-specific resolution (38-42).

Interestingly, for Tar32, its addition at ratios <0.25 triggered no significant shift of HSQC peaks of WT PLD (Supplementary Figure 3). However, at the ratio of 1:0.25 where the turbidity has the highest value and many droplets were formed, a large set of HSQC peaks became too broad to be detected, which appeared to result from the formation of large and dynamic Tar32-PLD complexes whose HSQC peaks became too broad to be detected. Nevertheless, with further addition of Tar32, most disappeared peaks were restored at 1:0.5 and no large shift was observed with the ratio further increased to 1:1, where all droplets were completely dissolved. Strikingly, at 1:1, a large set of HSQC peaks were significantly shifted as compared with those in the free state, indicating that a large set of residues were perturbed by Tar32.

We then titrated WT PLD with A6 under the same conditions. Briefly, A6 induced the broadening of HSQC peaks at 1:3 where the turbidity value reached the highest and many droplets were formed. The intensity of most broadened peaks became largely restored and the shift of HSQC peaks was mostly saturated at 1:5, where all droplets were dissolved (Supplementary Figure 4). Different from what was observed on Tar32, addition of A6 only triggered the shift of a small set of HSQC peaks whose pattern is very similar to that induced by ATP as we previously observed (37). Moreover, the titrations of A32 led to the changes of HSQC peaks with the overall pattern very similar to that by Tar32 (Supplementary Figure 5).

Detailed NMR assignments revealed that for A6, except for the residues Arg268 which had a significant shift by ATP but became too broad to be detectable by A6, as well as Met414 which had significant shift by ATP but no large shift by A6, the residues with significantly shifted HSQC peaks are very similar to those induced by ATP, which are all clustered over N-terminal three Arg residues including: Gln269-L270, Arg272-Ser273 and Arg275-Phe276 (Fig 1d). Strikingly, the peaks from these residues gradually shifted upon stepwise addition of A6 and the shift was largely saturated at 1:5 (Fig 1e). This indicates that the binding affinity of A6 is relatively weak and in the fast exchange regime of NMR chemical shift time scale (40).

By contrast, the residues significantly perturbed by Tar32 and A32 are much more profound and located over the whole sequence which include Glu271-Ser273, Arg275-Phe276, Gln286, Arg293-Gly294, Met307, Ala324, Ala328, Gln331, Leu340, Arg361, Gly368, Ile383, Ser407-Lys408 (Fig 1f). Overall, the shift patterns induced by Tar32 and A32 are very similar although the amplitude of shifts is different to some degree (Fig 1f). The results suggest that in binding with TDP-43 PLD: 1) Tar32 and A32 have the binding affinity much higher than that of A6, and thus their affinity is in the slow exchange regime of the NMR chemical shift tine scale (40); 2) Tar32 and A32 have the highly similar binding sites and affinity. As Tar32 and A32 have the same backbone but different bases, the NMR chemical shift changes together with the above turbidity and DIC results imply that different bases have the highly similar binding affinity to Arg/Lys residues.

The current results together with the previous ones with ATP (37) reveal that ATP, A6, Tar32 and A32 appear to bind Arg/Lys residues of TDP-43 PLD with a common mechanism as they are all composed of the same building unit: nucleotide. As such, they are all able to establish electrostatic interactions between phosphate group of nucleotide and side chain cations of Arg/Lys as well as π-π/π-cation interactions between base aromatic rings and Arg/Lys side chains. The results that Tar32 containing all four bases and A32 consist of only adenine have the highly similar modulating capacity suggest that four bases, regardless of adenine (A)/guanine (G) with a purine aromatic ring, or thymine (T)/cytosine (C) with a pyrimidine aromatic ring, all have a highly similar affinity in establishing π-π/π-cation interactions with side chains of Arg/Lys residues of IDRs.

Nevertheless, as ATP, A6, Tar32/A32 have different numbers of covalently-linked nucleotides, they are expected to have length-dependent affinities, because it is well known that for a multivalent binder, its dissociation constant (Kd) value is the time of Kd values of the individual binding events if assuming these binding events are independent (43). As such, ATP can only establish bivalent binding and therefore has a low affinity to Arg/Lys. On the other hand, A6 and A32/Tar32 have multiple covalently-linked nucleotides and thus can establish the multivalent binding. In particular, as A32/Tar32 is much longer than A6 and consequently one A6 molecule is only able to maximally achieve multivalent binding to the clustered Arg268, Arg272 and Arg275 but one Tar32/A32 molecule is able to bind most, if not all, Arg/Lys residues of TDP-43 PLD as evidenced by the NMR results that Tar32/A32 could perturb residues over the whole PLD molecule. As a result, Tar32/A32 bind PLD with an affinity much higher than that of A6. The perturbations to residues other than Arg/Lys residues by Tar32/A32 might not be due to the direct binding with A32/Tar32, but result from the avoidable close contacts of these residues with A32/Tar32, or/and conformational/dynamic changes of these residues induced by the binding of A32/Tar32 to Arg/Lys, as we previously observed on the binding of ATP with TDP-43 PLD (37). Indeed, previously we showed that ssDNAs showed no detectable binding to the 165-residue PLD of FUS, which is completely absent of Arg/Lys (38).

### ssDNAs bind Arg and Lys residues with distinctive affinities

Previously we found that the binding affinity of ATP to Arg is much higher that that to Lys (37). Here we addressed the question whether this observation also holds for ssDNAs by titrating A6, Tar32 and A32 into AllK-PLD with its pI (9.6) only slightly lower than that of WT-PLD (Fig 2a), which we previously constructed and characterized by replacing all five Arg residues with Lys (37). Intriguingly, previously we found that Arg residues in fact behave to conformation-specifically inhibit LLPS of TDP-43 PLD and consequently AllK-PLD could weakly undergo LLPS at 15 μM even in the free state, with a turbidity of 0.23 (Fig. 2a) and formation of some small droplets with diameters of ∼0.7 μm (Supplementary Figure 6).

**Fig 2.**
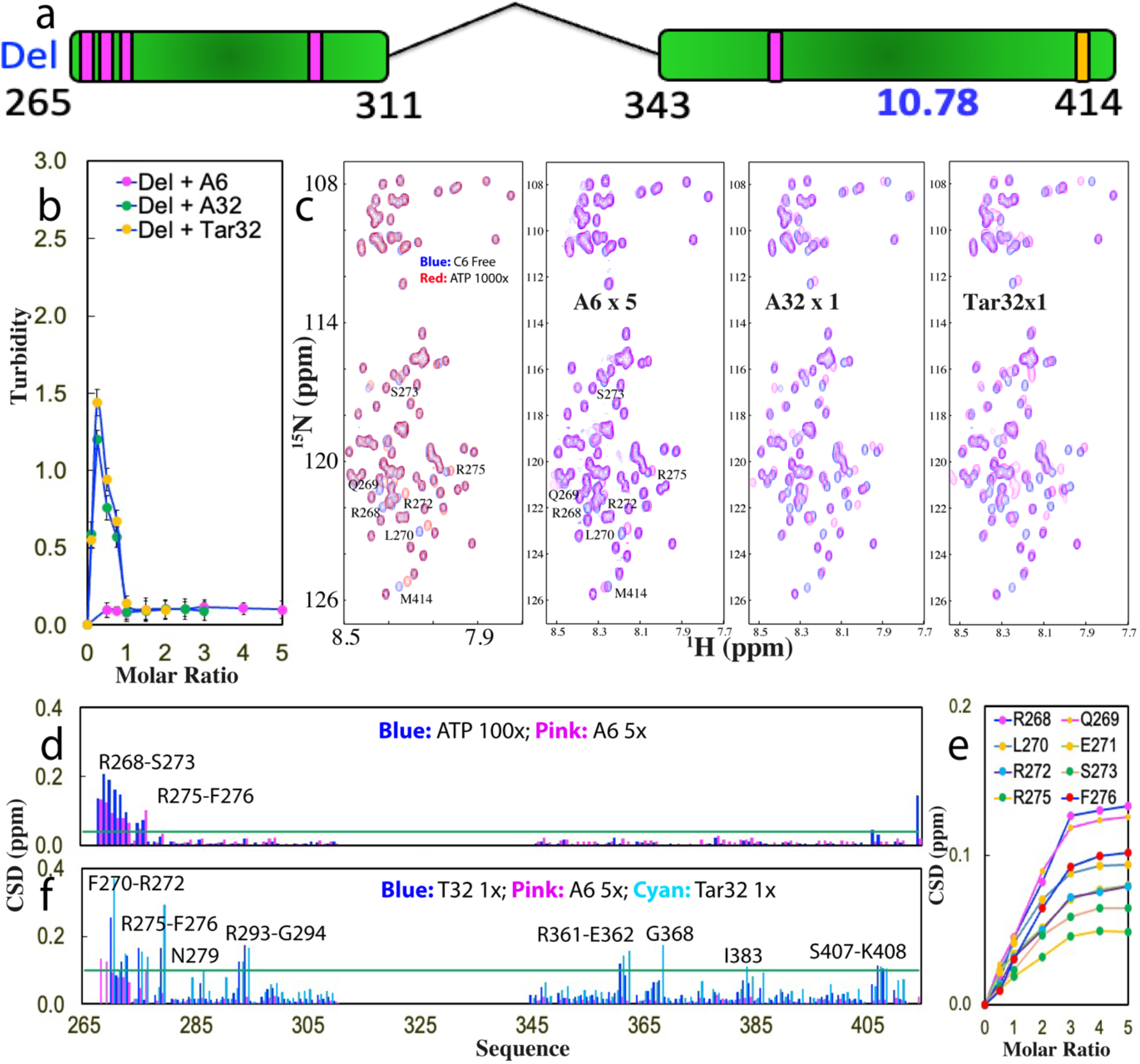
The hydrophobic region is not essential for LLPS driven by ssDNA. (a) Schematic representation of TDP-43 Del-PLD with residues 311-343 deleted. (b) Turbidity curves of Del-PLD in the presence of Tar32, A6 and A32 at different molar ratios. (c) HSQC spectra of Del-PLD in the absence (blue) and in the presence of ATP at a molar ratio of 1:1000 (red), as well as A6 at 1:5, A32 and Tar32 at 1:1 (purple). (d) Chemical shift difference (CSD) of Del-PLD between the free state and in the presence of ATP at 1:1000 (blue) and A6 at 1:5 (purple) respectively. The green line is used to indicate the value (0.04): Average + STD at the ratio of 1:5 (Del-PLD:A6). The residues with CSD values > Average + SD are defined as significantly perturbed residues and labeled. (e) CSD tracings of eight significantly perturbed residues in the presence of A6 at different ratios. (f) Chemical shift difference (CSD) of Del-PLD between the free state and in the presence A6 at 1:5 (purple), A32 at 1:1 (blue) and Tar32 at 1:1 (cyan) respectively. The green line is used to indicate the value (0.1): Average + STD in the presence Tar32 at the ratio of 1:1 (Del-PLD:Tar32). The residues with CSD values > Average + SD are defined as significantly perturbed residues and labeled.

We then titrated A6 into AllK-PLD but even up to 1:5, A6 triggered no significant change of turbidity (Fig. 2a) as well as number and size of droplets. By contrast, addition of Tar32 increased the turbidity which reached the highest (1.59) at 1:0.5. DIC imaging showed that a large amount of droplets were formed with some having diameter to ∼1.5 μm. Further increase of Tar32 concentrations led to the reduction of the turbidity and number of droplets and at 1:1, the turbidity was only 0.15 and no droplet could be observed. Titrations of A32 resulted in the similar turbidity curve as well as induction and dissolution of droplets.

Interestingly, HSQC titrations showed that addition of A6 had almost no large perturbation of HSQC peaks (Supplementary Figure 7) and even at 1:5, only two peaks from Lys268 and Met414 showed small shifts, thus suggesting that the binding affinity of A6 to AllK-PLD is very weak. On the other hand, addition of Tar32 triggered no large shifts of HSQC peaks of AllK-PLD at 1:0.1 but induced dramatic peak broadening at 1:0.5 (Supplementary Figure 8), where the turbidity reached the highest. Further addition of Tar32 at 1:0.75 led to restore of some disappeared peaks but many HSQC peaks still remained too broad to be detectable. Intriguingly, at 1:1 where the droplets were completely dissolved, many peaks detectable at 1:0.75 became too broad to be detectable.

A very similar pattern of HSQC peak changes was observed on the titrations by A32 (Supplementary Figure 9). The results imply that the binding affinity of Tar32/A32 to AllK-PLD is higher than that of A6 to WT-PLD but weaker than that of Tar32/A32 to WT-PLD. As such, the binding affinity of Tar32/A32 to AllK-PLD is in the intermediate exchange regime of the NMR chemical shift time scale characteristic of severe broadening of NMR peals, while that of A6 to WT-PLD is in the fast exchange regime and that of Tar32/A32 to WT-PLD is in the slow exchange regime.

Therefore, unlike A6 whose addition induced a gradual shift of HSQC peaks of WT-PLD and Tar32/A32 whose addition resulted in two sets of HSQC peaks of WT-PLD, the addition of Tar32/A32 into AllK-PLD even at an exceeding amount resulted in extensive broadening/disappearance of HSQC peak. The results together thus indicate that like what we observed with ATP (37), the binding affinity of ssDNAs to Arg residues is also much higher than that to Lys, consistent with the previous observation on RNA (41), because Arg side chain with a planar and delocalized guanidinium cation can establish both π-π and π-cation interactions with base aromatic ring, while Lys side chain with a tetrahedral ammonium cation can only establish π-cation interactions with base aromatic ring (Supplementary Figure 1).

### Nucleic-acid-driven mechanism of LLPS of TDP-43 PLD

The above results reveal that like ATP, three ssDNAs are able to biphasically modulate LLPS of RDP-43 WT-PLD by the specific binding to Arg/Lys. We then aimed to understand the nucleic-acid-driven mechanism of LLPS of TDP-43 PLD. Previously, in the absence of nucleic acids, the key driving force for LLPS of PLD has been extensively characterized to result from the dimerization/oligomerization of the hydrophobic region 311-343 uniquely existing in TDP-43 PLD. Furthermore, we have previously shown that for Del-PLD with the residues 311-343 deleted (Fig 3a), ATP no longer induced its phase separation although it still binds the same set of residues with the highly similar affinity, indicating the intrinsic driving force is essential for ATP to induce LLPS (37).

**Fig 3.**
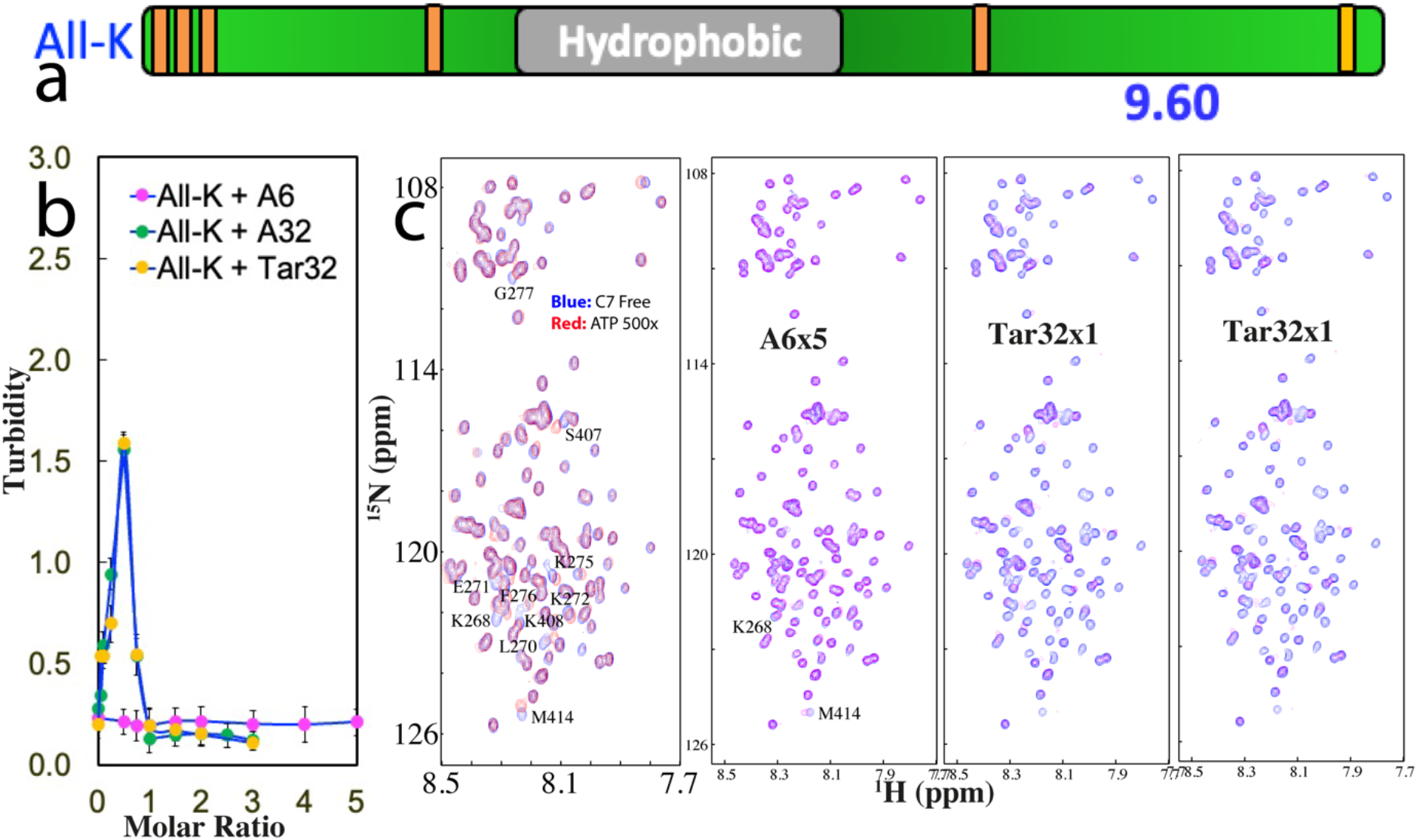
Three ssDNAs bind to Arg with the affinity higher than that to Lys. (a) Schematic representation of AllK-PLD with all five Arg mutated to Lys. (b) Turbidity curves of AllK-PLD in the presence of Tar32, A6 and A32 at different molar ratios. (c) HSQC spectra of AllK-PLD in the absence (blue) and in the presence of ATP at a molar ratio of 1:500 (red), as well as A6 at 1:5, A32 and Tar32 at 1:1 (purple) respectively.

We first titrated Del-PLD with A6 but no significant change was observed on the turbidity with the ratios even up to 1:5 (Fig 3b). DIC imaging confirmed that no droplet was formed with the ratios up to 1:5. The results indicate that A6 was unable to induce LLPS of Del PLD. By contrast, upon titration by Tar32, the turbidity increased rapidly and reached the highest value (1.44) at 1:0.25 (Fig 3b). DIC imaging showed that many droplets were observed with the diameters of some droplets reaching ∼1.5 μm (Supplementary Figure 10). Further addition led to the reduction of turbidity and at 1:1 the turbidity is only 0.14 where no droplets were detected by DIC imaging. Interestingly, the titrations with A32 resulted in the patterns of turbidity and DIC changes very similar to those by Tar32.

We also characterized the interaction of Del-PLD with A6, Tar32 and A32 by NMR HSQC titrations (Supplementary Figures 11-13). Interestingly, although A6 is incapable of inducing LLPS of Del-PLD, it could still trigger the shift of a small set of HSQC peaks very similar to that of Del-PLD induced by ATP, which include Arg268-Ser273 and Arg275-Phe276 (Fig 3d). Similar to what was observed on WT-PLD titrated by A6 (Fig 1e), HSQC peaks of Del-PLD residues also gradually shifted upon stepwise addition of A6 and the shift was largely saturated at 1:5 (Fig 3e). By contrast, the residues significantly perturbed by Tar32 and A32 are much more profound which are over the whole sequence which include Phe270-Arg272, Arg275-Phe276, Asn279, Arg293-Gly294, Arg361-Glu362, Gly368, Ile383, Ser407-Lys408 (Fig 3f).

The results indicate that despite deletion of residues 311-343, the binding affinity of A6 to Del-PLD is very similar to that of WT-PLD and also in the fast exchange regime, while those of Tar32/A32 remain in the slow exchange regime. Therefore, the existence of the hydrophobic region 311-343 appears to have no detectable impact on the binding affinity of A6, A32 and Tar32 to Arg/Lys. Nevertheless, it has the important contribution to the strength of the driving force for LLPS. For A6 with the low binding affinity, although it still binds the similar set of residues of Del-PLD at similar affinity, it failed to induce LLPD because of the absence of the intrinsic driving force from the oligomerization of residues 311-343. Nevertheless, for Tar32/A32 with a strong binding affinity, their multivalent binding is sufficient to drive and then dissolve LLPS of Del-PLD even without the intrinsic driving force. So it becomes clear that although for the special case of TDP-43 WT-PLD, the intrinsic driving force is essential for driving LLPS in the absence of nucleic acids, a nucleic acid with sufficient length and binding affinity is sufficient to drive LLPS by multivalently and specifically binding Arg/Lys residues even with the intrinsic driving force completely deleted.

### ATP and ssDNAs interplay to modulate LLPS by competing for binding Arg/Lys

As ATP and nucleic acids have been shown to specifically bind Arg/Lys residues, we thus set to assess whether ATP and nucleic acids interplay in modulating LLPS of TDP-43 PLD by titrating ATP into four phase separated samples: namely WT-PLD in the presence of A6 at 1:3 (Supplementary Figure 14); WT-PLD in the presence of Tar32 at 1:0.25 (Supplementary Figure 15); AllK-PLD in the presence of Tar32 at 1:0.5 (Supplementary Figure 16); and Del-PLD in the presence of Tar32 at 1:0.25 (Fig 4).

**Fig 4.**
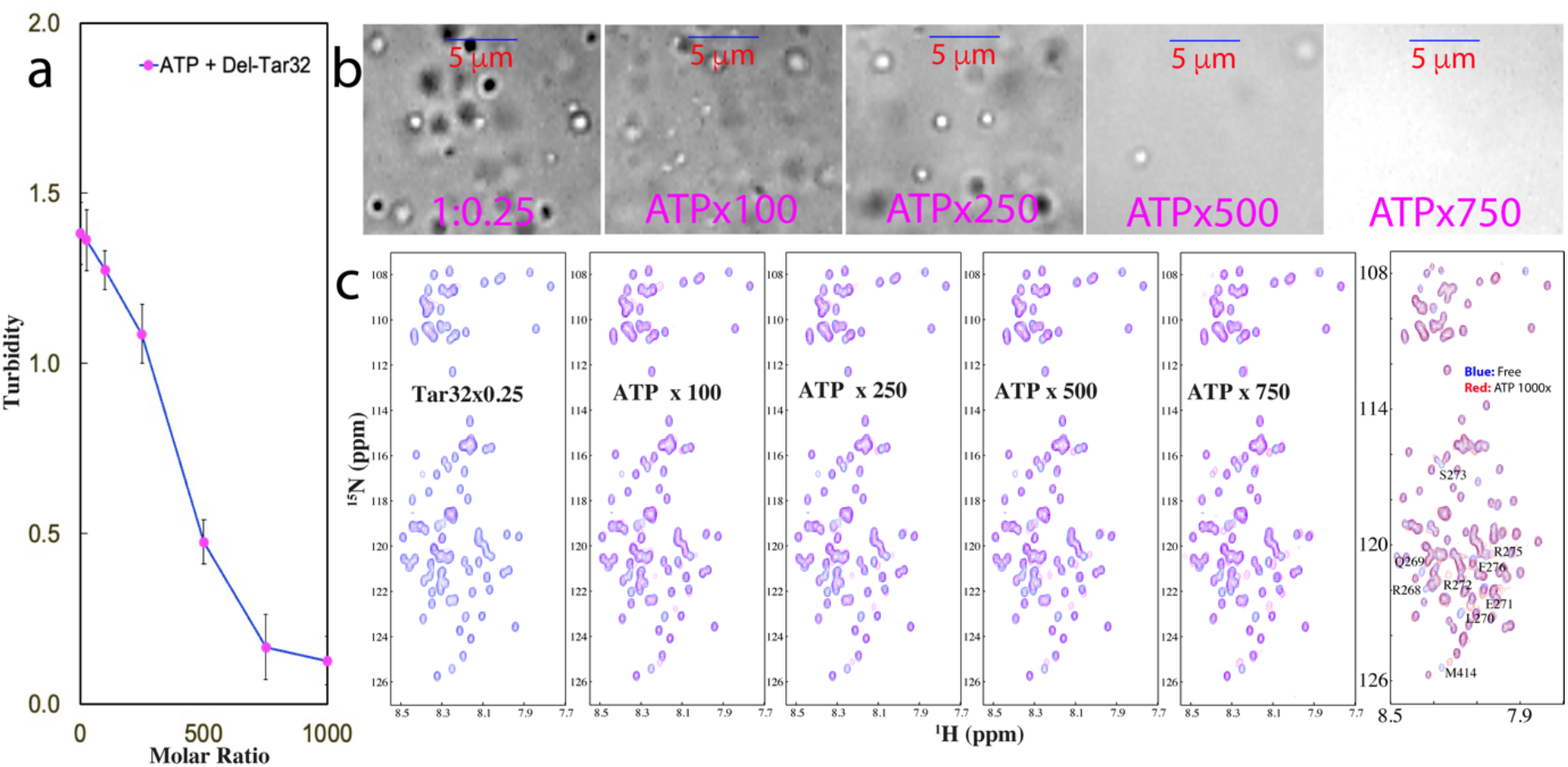
ATP dissolves LLPS of Del-PLD maintained by Tar32. (a) Turbidity curve of Del-PLD in the presence of Tar32 at 1:0.25, with further addition of ATP at different molar ratios. (b) DIC images of Del-PLD in the presence of Tar32 at 1:0.25, and with further addition of ATP at different molar ratios. (c) HSQC spectra of Del-PLD in the absence (blue) and in the presence of Tar32 at 1:0.25 (purple), as well as with further addition of ATP at different molar ratios.

Very unexpectedly, for WT-PLD in the presence of A6 at 1:3, the addition of ATP to 1:25 completely dissolved the liquid droplets as evidenced by the reduction of turbidity to 0.12, as well as dissolution of all droplets by DIC imaging. Strikingly, NMR HSQC titration indicates that upon addition of ATP even at 1:25, the intensity of many broadened HSQC peaks due to phase separation in the presence of A6 become largely restored. Nevertheless, a number of peaks still remained to be very broad (Supplementary Figure 14). Intriguingly, further addition of ATP to 1:50 rendered some restored peaks to become too broad to be undetectable and more peaks became undetected at 1:100. This may be due to the occurrence of dynamic aggregation induced by the non-specific screening effect of ATP and A6, both of which are highly negatively charged. Further addition of ATP above 1:100 resulted in a complete precipitation of WT PLD and consequently no NMR signals could be detected.

For WT PLD in the presence of Tar32 at 1:0.25, addition of ATP up to 1:750 resulted in no significant changes in turbidity and droplet number/size, implying that ATP exerts minor effect on disrupting the Tar32-PLD complex which is jointly driven by both strong nucleic-acid-binding and intrinsic driving force. As monitored by NMR, addition of ATP not only failed to restore the disappeared HSQC peaks, but rendered more peaks to become broad and at 1:1000, most HSQC peaks became too broad to be detectable (Supplementary Figure 15). This implies that the competition between ATP and Tar32 for binding Arg/Lys residues may shift the binding affinity of Tar32 to WT-PLD from the slow exchange regime to intermediate exchange regime, or dynamic aggregation occurred to some degree, both of which lead to severe broadening of HSQC peaks. Intriguingly, further addition of ATP above 1:750 resulted in a complete precipitation and no NMR signal could be detected.

For AllK-PLD in the presence of Tar32 at 1:0.5, addition of ATP up to 1:250 only resulted in a slight reduction in the number of the droplets (Supplementary Figure 16a). On the other hand, as monitored by NMR, addition of ATP to 1:50 led to the appearance of some peaks but further addition resulted in further disappearance of HSQC peaks. Upon addition of ATP above 1:250, the sample was completely precipitated and consequently no NMR signal could be detected.

Interestingly, for Del-PLD in the presence of Tar32 at 1:0.25, addition of ATP led to the continuous dissolution of LLPS as indicated by the reduction of turbidity (Fig 4a) and dissolution of droplets. At 1:750, all droplets have been completely dissolved (Fig 4b) and no aggregation was observed even up to 1:1000. Most interestingly as shown in Fig 4c, at 1:100, the majority of disappeared NMR peaks due to the presence of Tar32 has been restored. Nevertheless, some NMR peaks still have large shifts as compared to those in the free state or only in the presence of ATP. Addition of ATP to 1:500 resulted in HSQC spectrum which is very similar to that only in the presence of ATP except for several peaks. Further addition of ATP to 1:1000 led to no further shifts.

In summary, the results together reveal that ATP and ssDNAs do interplay in modulating LLPS by competing for binding Arg/Lys. For the phase separated state of Del-PLD with the hydrophobic region deleted which is thus only driven by the nucleic-acid-binding to Tar32, ATP is capable of completely dissolving it without inducing any aggregation. However, For the phase separated states of WT-PLD in the presence of A6 or Tar32 as well as AllK-PLD in the presence of Tar32, which are maintained by both nucleic-acid-binding and oligomerization of the hydrophobic region, the competition between ATP and nucleic acids for binding Arg/Lys could all lead to exaggeration of LLPS into aggregation. Here we propose a potential mechanism: because ATP and nucleic acids are highly negatively charged, they can exert at least two strong electrostatic effects: namely site-/conformation-specific electrostatic interaction and non-specific screening effect. If ATP and nucleic acids are relatively tightly bound with Arg/Lys residues, site-/conformation-specific negative charges will be introduced onto PLD molecules. Consequently the site-/conformation-specific repulsive electrostatic interaction will become dominant which not only contributes to dissolving LLPS, but also to preventing the aggregation triggered by hydrophobic interaction. By contrast, If they fail to be relatively tightly bound with Arg/Lys residues, the non-specific screening effect will become dominantly operating which will trigger aggregation driven by hydrophobic interaction, as we universally found on the partially-folded or disordered proteins with significant exposure of hydrophobic patches, which are all soluble in unsalted water but become insoluble upon the introduction of salts even at very low concentrations (44).

## Discussion

Although most, if not all, MLOs contain both nucleic acids and IDR-rich proteins, currently nucleic acids have been largely considered to exert non-specific electrostatic/salt effects onto LLPS. The present study, however, deciphers that ssDNAs are capable of driving and then dissolving LLPS of TDP-43 PLD with the capacity dependent on the length but not sequence of ssDNAs. Very unexpectedly, extensive NMR characterization unambiguously provides the first residue-resolution evidence that ssDNAs achieve the modulation of LLPS mainly by the multivalent binding specifically to Arg/Lys residues with the affinity to Arg higher than that to Lys.

Together with recent NMR results with ATP (37,39,45,46), a common mechanism is emerging for ATP and nucleic acids to modulate LLPS of IDRs by specific binding. Briefly, different from the binding to a well-folded protein to form the stable classic complex with a well-defined three-dimensional structure in which various types of residues are involved and a single atom variation of ATP/nucleic acids or proteins might dramatically alters the binding affinity, in forming the dynamic complex with IDRs which lack the defined conformation and thus are highly accessible to the bulk solvent, ATP/nucleic acids can only establish the NMR-detectable binding to Arg/Lys residues through electrostatic interactions between phosphate groups of ATP/nucleic acids and side chain cations of Arg/Lys as well as π-π/π-cation interactions between base aromatic rings and Arg/Lys side chains. In particular, different bases with distinctive aromatic rings have a highly similar affinity to Arg/Lys residues of IDRs, indicating that the binding specificity for forming the dynamic nucleic-acid-IDR complexes is lower than that for the classic nucleic-acid-protein complexes. In this context, RNA and ssDNA are expected to bind Arg/Lys residues of IDRs with the same mechanism, because they only have two minor differences in their chemical structures.

On the other hand, nucleic acids differ from ATP in having multiple covalently-linked nucleotides and consequently are capable of establishing multivalent binding to Arg/Lys to gain the length-dependent affinity. Furthermore, because ATP and nucleic acids are highly negatively charged, they thus own the capacity to exert strong electrostatic effects onto LLPS and aggregation of IDRs, which, however, appear to be context-dependent. For example, ATP and nucleic acids impose at least dual effects on LLPS and aggregation of TDP-43 WT-PLD. Briefly, if in the presence of exceeding amounts, ATP or nucleic acids are able to become relatively tightly bound with its Arg/Lys residues, PLD molecules will become site-/conformation-specifically associated with multiple negative charges, whose repulsive electrostatic interaction acts not only to disrupt LLPS, but also to prevent its irreversible aggregation. By contrast, if ATP or/and nucleic acids are unable to be bound with PLD, their non-specific screening effect will become dominantly operating to trigger aggregation driven by oligomerization of the unique hydrophobic region, as we universally observed on various aggregation-prone or even “insoluble” proteins with exposed hydrophobic patches, which are all soluble in salt-free water but become aggregated in salted solution (44).

In this framework, the mechanisms for three ssDNAs to interact with three PLDs can be formulated. As illustrated in Fig 5, the short A6 appears to be only able to maximally achieve multivalent binding to the clustered Arg268, Arg272 and Arg275 and gains a relatively weak affinity. Therefore, only in the presence of the intrinsic driving force from the oligomerization of the hydrophobic region, A6 is able to drive LLPS of WT-PLD to form dynamically and multivalently cross-linked A6-PLD complex which manifests as liquid droplets. Further addition of A6 to an exceeding amount will lead to the formation of A6-boumd PLD which is highly negatively-charged. Consequent the droplets will be disrupted as well as the aggregation of WT-PLD triggered by the oligomerization of the hydrophobic region is inhibited by the repulsive electrostatic interaction (I of Fig 5a). In this regard, despite binding to the same Arg residues of Del-PLD with a similar affinity, A6 can no longer drive its LLPS because of the deletion of the intrinsic driving force (II of Fig 5a). Furthermore, due to the much weaker binding affinity between the base ring and Lys side chain, A6 has almost no detectable binding to Lys of AllK-PLD as well as is unable to drive its LLPS (III of Fig 5a). By contrast, in addition to binding the clustered Arg268, Arg272 and Arg275, the long Tar32/A32 appears to further bind Arg293, Arg361 and Lys408 and consequently acquire the affinity much high than that of A6. As such, for WT-PLD, Tar32/A32 drives LLPS by coupling both nucleic-acid-binding and intrinsic driving force to form the dynamic and multivalent Tar32/A32-PLD complex also manifesting as liquid droplets. Similarly, further addition of Tar32/A32 will also lead to the formation of highly negatively-charged Tar32/A32-boumd PLD. Consequently, the droplets will be disrupted and the aggregation is inhibited (I of Fig 5b). Importantly, due to the high binding affinity, Tar32/A32 is still able to drive and then dissolve LLPS of the 117-residue Del-PLD even with the intrinsic driving force deleted (II of Fig 5b). Furthermore, although the individual binding affinity of each base ring to Lys is much weaker than that to Arg, the overall affinity resulting from the multivalent binding of long Tar32/A32 to multiple Lys residues of AllK-PLD is still sufficiently high to drive and then dissolve its LLPS (III of Fig 5b).

**Fig 5.**
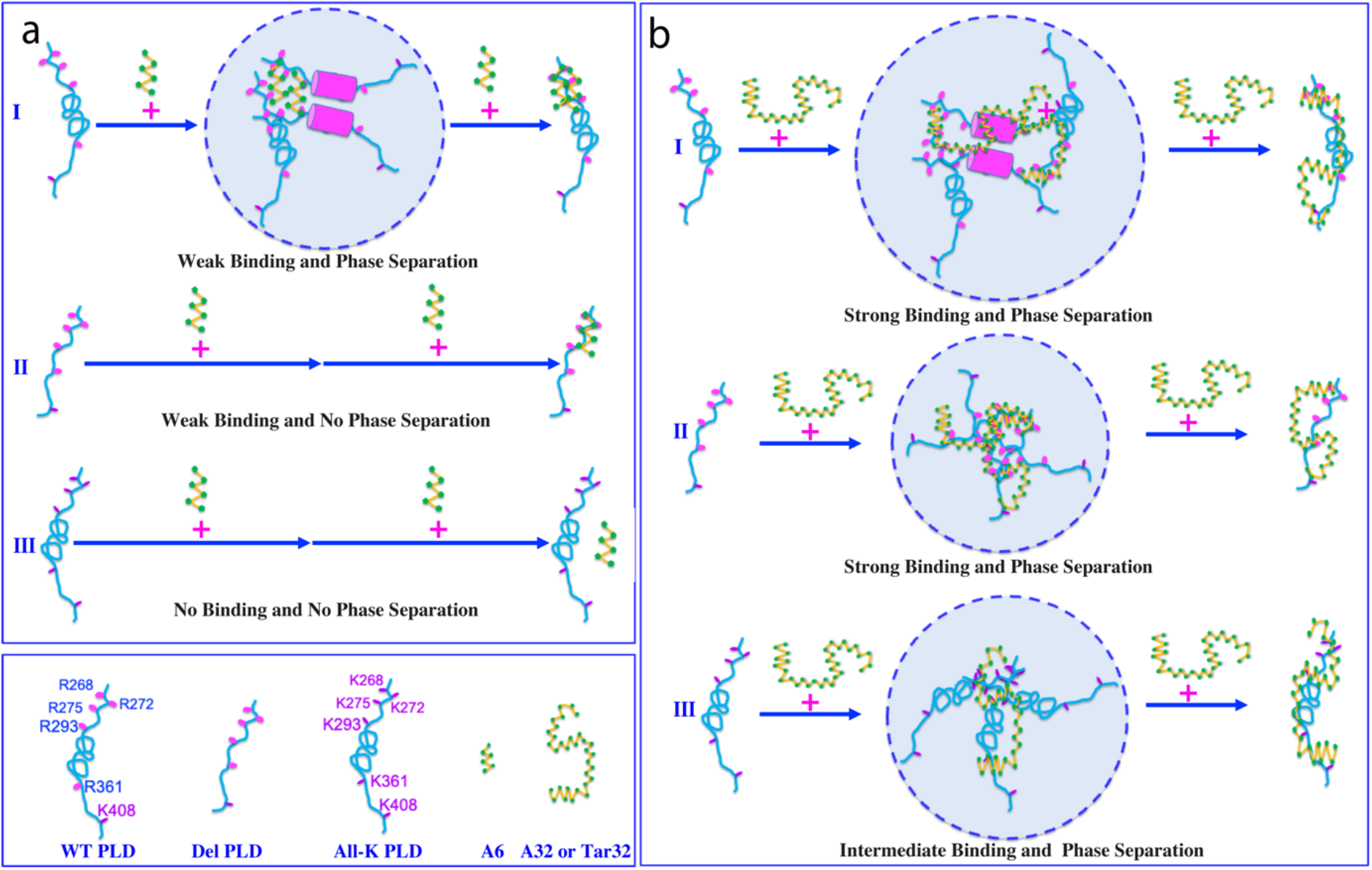
Speculative mechanisms to illustrate interactions of three ssDNAs to three PLDs. (a) Interactions of A6 to three PLDs. (b) Interactions of Tar32/A32 to three PLDs.

These results together not only unveil the first nucleic-acid-driven mechanism for LLPS of TDP-43 PLD; but also decode a general principle that not just TDP-43 PLD, all Arg/Lys-containing IDRs, which are not limited to those rich in RG-/RGG-motifs as the C-terminal domain of FUS (38), can act as ATP and nucleic-acid-binding domains. Most strikingly, the multivalent and specific binding of their Arg/Lys residues with nucleic acids of a sufficient length is sufficient to drive and dissolve their LLPS without the need of any other driving force. Furthermore, ATP and nucleic acids universally interplay to modulate LLPS by competing for binding Arg/Lys of IDRs. However, for TDP-43 WT-PLD containing the hydrophobic region, whose oligomerization also serves to drive LLPS, its LLPS is prone to exaggeration into irreversible aggregation, particularly when ATP and nucleic acids co-exist to compete for binding Arg/Lys. Their competition may lead to reduction of the binding affinity of both of them. Consequently, the negative charges of ATP or/and nuclei acids site-/conformation-specifically associated with Arg/Lys of PLD will become dissociated to some degree. Under such a circumstance, the repulsive electrostatic interaction among PLD molecules will be reduced while the non-specific screening effects imposed by ATP and nucleic acids will become operating dominantly to enhance oligomerization of the hydrophobic region, which eventually results in exaggerating LLPS into the irreversible aggregation.

## Conclusion

In contrast to the wide belief, the present study decodes for the first time that mainly by multivalently and specifically binding Arg/Lys residues, nucleic acids act to drive and then dissolve LLPS of TDP-43 PLD with the length-dependent capacity. Although for the special case of TDP-43 WT-PLD, the nucleic-acid-binding drives its LLPS by coupling the intrinsic driving force, in the general context the multivalent nucleic-acid-binding itself is sufficient to drive and then dissolve LLPS of Arg-/Lys-containing IDRs without the need of any other driving force. Most strikingly, ATP and nucleic acids have been decrypted to share the common mechanism in biding Arg/Lys of IDRs. Consequently, not just TDP-43 PLD, all Arg/Lys-containing IDRs are cryptic domains for binding ATP and nucleic acids and may thus phase separate upon multivalently binding nucleic acids.

Most strikingly, ATP and nucleic acids universally interplay in modulating LLPS by competing for binding Arg-/Lys. This discovery bears unprecedented implications in our understanding of the homeostasis of fundamental cellular processes. So far, even RG-/RGG-rich IDRs have been identified in >1,700 human proteins (47-51). Generally, it was estimated that the average frequency of Arg found in eukaryotic protein is ∼5.6% while that of Lys is ∼6.2 (52). As such, IDRs are anticipated to contain multiple Arg/Lys whose number is comparable to or even higher than that of TDP-43 PLD unless their frequency of Arg/Lys is strongly biased and lower than the average frequency. Indeed, for example, the 40-residue intrinsically-disordered Nogo-40, which has no known function to bind ATP and nucleic acids to undergo LLPS, contains two Arg and three Lys residues (53). In this context, most, if not all, IDRs, which account for about the half of human proteome, are capable of multivalently binding nucleic acids to undergo LLPS. Nevertheless, the abnormally high affinity of Arg/Lys-containing IDRs to nucleic acids may unavoidably provoke cytotoxicity. For example, the ALS-causing C9orf72 dipeptide repeats extremely rich in Arg have been recently shown to have an extremely high affinity and consequently to generally displace RNA/DNA-binding proteins from binding mRNA in chromatin, which consequently impairs any processes involving nucleic acids (54).

Intriguingly, only a very limited number of MLOs have been identified in cells. One possible mechanism underlying this puzzling observation is that ATP with high cellular concentrations might play a previously-unrecognized role in inhibiting LLPS of most of Arg/Lys-containing IDRs. It appears that even for the formed IDR-nucleic-acid complexes and MLOs such as SGs, ATP is still essential to maintain their functional dynamics or reversibility as well as to prevent their exaggeration into irreversible aggregation, which are associated with an increasing spectrum of human diseases including all neurodegenerative diseases.

## Methods

### Preparation of recombinant WT and mutated TDP-43 PLD proteins

Here, we used our previously-cloned DNA constructs in a modified vector without any tag which include those encoding TDP-43 WT-PLD over residues 265-414, Del-PLD with residues 311-343 deleted and AllK-PLD with all five Arg replaced by Lys (37). All three recombinant TDP-43 PLD proteins were highly expressed in *E. coli* BL21 cells and were found in inclusions. Consequently, they were purified by the previously established protocols in other and our labs. Briefly the recombinant proteins were solubilized with the buffer with 8 M urea and the Reverse Phase (RP)-HPLC purification was used to obtain highly pure proteins with the impurities including liquid, ions and nucleic acids removed (37). Isotope-labeled proteins for NMR studies were prepared by the same procedures except that the bacteria were grown in M9 medium with addition of (^15^NH_4_)_2_SO_4_ for ^15^N-labeling. The protein concentration was determined by the UV spectroscopic method in the presence of 8 M urea, under which the extinct coefficient at 280 nm of a protein can be calculated by adding up the contribution of Trp, Tyr, and Cys residues (37,55).

ATP were purchased from SigmaAldrich with the same catalog numbers as previously reported (37). Three synthetic ssDNAs were purchased from a local company (33). Proteins, ssDNAs and ATP samples were all prepared in 10 mM sodium phosphate buffer and MgCl_2_ at the equal molar concentration to ATP was added for stabilization by forming the ATP-Mg complex (37). The final solution pH values were checked by pH meter and the small variations were adjusted with aliquots of very diluted NaOH or HCl (37).

### Differential interference contrast (DIC) microscopy and turbidity measurement

The formation of liquid droplets was imaged at 25 °C on 50 µl of different TDP-43 PLD samples at 15 μM in 10 mM sodium phosphate in the absence and in the presence of ssDNAs at different molar ratios by diffe rential interference contrast (DIC) microscopy (OLYMPUS IX73 Inverted Microscope System with OLYMPUS DP74 Color Camera) (37). The turbidity measurement and DIC imaging were performed after 15 min of the sample preparation. The turbidity was measured three times at wavelength of 600 nm and reported as Average + SD.

### NMR characterizations

All NMR experiments were acquired at 25 °C on an 800 MHz Bruker Avance spectrometer equipped with pulse field gradient units and a shielded cryoprobe (37). To have the enhancing effect of the cryoprobe for NMR signal sensitivity, which is essential for NMR HSQC titration experiments at such a low protein concentration (15 µM), NMR samples had to be prepared in 10 mM sodium phosphate buffer, while pH value was optimized to 5.5 as many HSQC peaks of TDP-43 PLD disappeared at higher pH values due to the enhanced exchange with bulk solvent and/or dynamic association (56,57).

For NMR titration studies of the interactions between TDP-43 WT/mutated PLD proteins and ATP and ssDNAs, one-dimensional proton and two dimensional ^1^H-^15^N NMR HSQC spectra were collected on ^15^N-labelled samples at a protein concentration of 15 µM in 10 mM sodium phosphate buffer (pH 5.5) at 25 °C in the absence and in the presence of ssDNAs or ATP at different molar ratios.

NMR data were processed by NMRPipe (58) and analyzed by NMRView (59). To calculate chemical shift difference (CSD) induced by interacting with ssDNAs, HSQC spectra were superimposed and subsequently the shifted peaks were identified, which were further assigned to the corresponding residues of TDP-43 PLD with the NMR resonance assignments previously achieved by us and other groups (37). The degree of the perturbation was reported by an integrated index calculated by the following formula (37,40):

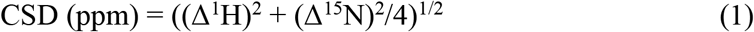

The residues with CSD values > Average + STD are defined as significantly perturbed residues.

## Acknowledgement

This study is supported by Ministry of Education of Singapore (MOE) Tier 1 Grant R-154-000-B92-114 to Jianxing Song. The funders had no role in study design, data collection and analysis, decision to publish, or preparation of the manuscript.

## Author Contributions

J.S. and M.D conceived and designed the experiments. M.D., T.L. S.Z. and J.S. performed the research, analyzed the data, J.S. wrote the manuscript.

